# Rapid hyperosmotic-induced Ca^2+^ responses in *Arabidopsis thaliana* exhibit sensory potentiation and establish involvement of plastidial KEA transporters

**DOI:** 10.1101/048330

**Authors:** Aaron B. Stephan, Hans-Henning Kunz, Eric Yang, Julian Schroeder

## Abstract

Plants experience hyperosmotic stress when faced with saline soils and possibly drought stress, but it is currently unclear how plants perceive this stress in an environment of dynamic water availabilities. Hyperosmotic stress induces a rapid rise in intracellular Ca^2^+ concentrations ([Ca^2+^]_i_) in plants, and this Ca^2+^ response may reflect the activities of osmo-sensory components. Here, we find in the reference plant *Arabidopsis thaliana* that the rapid hyperosmotic-induced Ca^2+^ response exhibited enhanced response magnitudes after pre-exposure to an intermediate hyperosmotic stress. We term this phenomenon “osmo-sensory potentiation”. The initial sensing and potentiation occurred in intact plants as well as in roots. Having established a quantitative understanding of WT responses, we investigated effects of pharmacological inhibitors and candidate channel/transporter mutants. Quintuple MSL channel mutants as well as double MCA channel mutants did not affect the response. However interestingly, double mutations in the plastid KEA transporters, *kea1kea2*, and a single mutation that does not visibly affect chloroplast structure, *kea3*, impaired the rapid hyperosmotic-induced Ca^2+^ responses. These mutations did not significantly affect sensory potentiation of the response. These findings suggest that plastids may play an important role in the early steps mediating the response to hyperosmotic stimuli. Together, these findings demonstrate that the plant osmosensory components necessary to generate rapid osmotic-induced Ca^2+^ responses remains responsive under varying osmolarities, endowing plants with the ability to perceive the dynamic intensities of water limitation imposed by osmotic stress.

**Significance Statement:** The sensitivity ranges of biological sensors determine when‐ and to what extent responses to environmental stimuli are activated. Plants may perceive water limitation imposed by soil salinity or drought in the form of osmotic stress, among other mechanisms. Rapid osmotic stress-induced Ca^2+^ responses provide the opportunity to quantitatively characterize the responses to osmotic stress under environmental and genetic perturbations. This report describes a phenomenon whereby prior exposure to osmotic stress increases the sensitivity of the rapid responses to subsequent stress. Further, mutations in specific plastidial transporters were found to reduce the stress response. These findings inform the reader of new avenues for understanding osmotic stress responses in plants.

## Introduction

Plants exhibit a wide range of physiological responses to cope with water deprivation by drought and salinity stress (1–3). The properties of biological sensors determine the circumstances and extent to which these coping mechanisms are activated, but the early sensory mechanisms and components regulating the osmotic sensory components in plants are not well understood (4, for review). Pioneering studies have demonstrated that *Arabidopsis* seedlings expressing the bioluminescent Ca^2+^ reporter protein aequorin exhibit a rapid rise in intracellular Ca^2+^ ( [Ca^2+^]_i_) within seconds upon stimulation by NaCl solution (5, 6). This rapid osmotic-induced Ca^2+^ response has been observed in plant species ranging from rice (7) to the basal-branching moss taxon *Physcomitrella patens* (8), indicating that this response may be conserved across the Plantae kingdom. Solutions of either NaCl or iso-osmotic mannitol/sorbitol induce nearly identical rapid Ca^2+^ responses, indicating that the nature of this rapid stimulus is largely osmotic rather than ionic (5, 9, 10). Individual seedling responses tend to be quite heterogeneous, and the response characteristics show variation depending on the accession or extracellular ionic composition (9, 11). It has been demonstrated that, for NaCl stress in particular, Ca^2+^ influx first appears in roots and later in shoots (9), and when roots are stimulated with salt stress, the Ca^2+^ signal propagates to the leaves in waves (12, 13). The root-to-shoot propagation requires the slow vacuolar (SV) channel TPC1, proposed to mediate a Ca^2+^-induced Ca^2+^ release mechanism (12). However, TPC1 is not required for the rapid osmotic induced Ca^2+^ response (12). On a longer time scale, Ca^2+^ oscillations have been observed within individual cells (14). Thus, rapid hyperosmotic-induced Ca^2+^ responses represent the initial cytosolic Ca^2+^ rise preceding these secondary Ca^2+^ signaling events that occur throughout the plant.

Given the short time interval between stimulus and response, quantitative interrogation of the rapid hyperosmotic-induced Ca^2+^ response in plants was used as a method to identify a potential osmosensory component (15), as well as sensory components for other stimuli such as ATP and RALF peptides (16, 17). One mutant line, *reduced hyperosmolality-induced [Ca*^*2*+^]_*i*_ *increase 1 (osca1)*, was mapped to mutations in a membrane protein At4g04340 (15). Osca1 and a homolog CSC1 (encoded by the gene At4G22120) were both shown to confer osmotically-sensitive Ca^2+^-permeable channel currents (15, 18). It remains unknown, however, how these and other unidentified components of the osmo-sensory components are regulated.

Osmotic stress in plants is dynamic in nature; availability of water to the plant can vary widely depending on soil water capacity, dissolved ionic content, precipitation, atmospheric humidity, temperature, wind speed and solar irradiance (19). However, it is not known whether a dynamic osmotic environment modulates the activity of the initial osmosensory components in plants. Here we present a quantitative study of rapid hyperosmotic-induced Ca^2+^ responses in the plant *Arabidopsis thaliana*. Under conditions where the seedlings were pre-exposed to mild osmotic stress, we found that seedlings exhibited larger, more robust rapid Ca^2+^ responses to subsequent hyperosmotic stress. We term this response “osmo-sensory potentiation”.

Ca^2+^ is not only released across the plasma membrane from the apoplast; several organelle types may act as intracellular Ca^2+^ storage pools (20, 21). We further investigated several candidate mutant genes and found that members of the plastidial KEA transporter family are necessary for eliciting wild-type levels of rapid hyperosmotic-induced Ca^2+^, but that *kea* mutant lines were still largely capable of exhibiting sensory potentiation. Together, these results demonstrate that the sensitivities of the osmo-sensory components are capable of being tuned according to previous stimulation and indicate an important role for plastidial ion transporters in stress-induced [Ca^2+^]_i_ elevations.

## Results

### Features of rapid hyperosmotic-induced Ca^2+^ responses

Wild-type *Arabidopsis* seedlings of the ecotype Col-0 expressing aequorin under control of the strong and constitutive 35S promoter (henceforth referred to as “wild-type Col-0”) were grown for one week in individual wells of 96-well plates. The seedlings were pre-treated with the co-factor coelenterazine to reconstitute the active aequorin Ca^2+^ reporter complex. Stimulation solutions were automatically applied to each well while Ca^2+^-dependent light emission from individual seedlings was measured. We observed that stimulation of the seedlings by hyperosmotic stress resulted in a rapid, transient rise in intracellular Ca^2+^ (Fig. 1A-D), consistent with previous reports (5, 9, 12, 15). When seedlings were stimulated by a low osmolarity solution iso-osmotic to the existing media, only small responses were observed (Fig. 1A-B). These small responses could be the result of the seedlings sensing a mechanical force caused during application of the stimulus into the wells containing the seedlings **(22, 23)**. We found that these “mechano-sensory” responses peaked consistently within two seconds and returned to baseline by 20 seconds under the imposed conditions (Fig 1A-B). As the osmolarities of the stimuli were increased, the amplitudes of the rapid Ca^2+^ responses increased dramatically (Figs. 1A-D). These osmosensory responses varied from seedling to seedling in terms of amplitudes and kinetics (Figure 1C-D), as has been noted previously (9). However, we observed that nearly all hyperosmotic-induced Ca^2+^ responses exhibited an initial peak in free cytosolic Ca^2+^ within 5 seconds of stress application, and they exhibited an additional peak or several secondary peaks that were much more variable in amplitudes and kinetics (Fig. 1C-D). Occasionally the secondary peaks blended in with the first peak, giving the impression of a single, larger peak (e.g., in Fig. 1C, black trace)(24). The variation in amplitude is not due to variation in aequorin expression levels, since relative Ca^2+^ concentrations were calculated based on calibrated normalization to total active aequorin protein in each seedling (25)(see Methods). Indeed, independent aequorin-transformant wild-type Col-0 lines varying by more than 5-fold in aequorin expression levels showed indistinguishable mean hyperosmotic-induced Ca^2+^ responses after normalization (Fig. S1). Due to the inherent variability from seedling-to-seedling, we developed an analysis program that quantifies over 40 features of individual seedling responses including primary and secondary peak amplitudes, time to peaks, decay rates, integrated areas under the peaks, and principal components, and automated plots measured features for individual seedlings as well as averages of replicate seedlings for any tested treatment or genotype (Materials and Methods).

**Figure 1.**
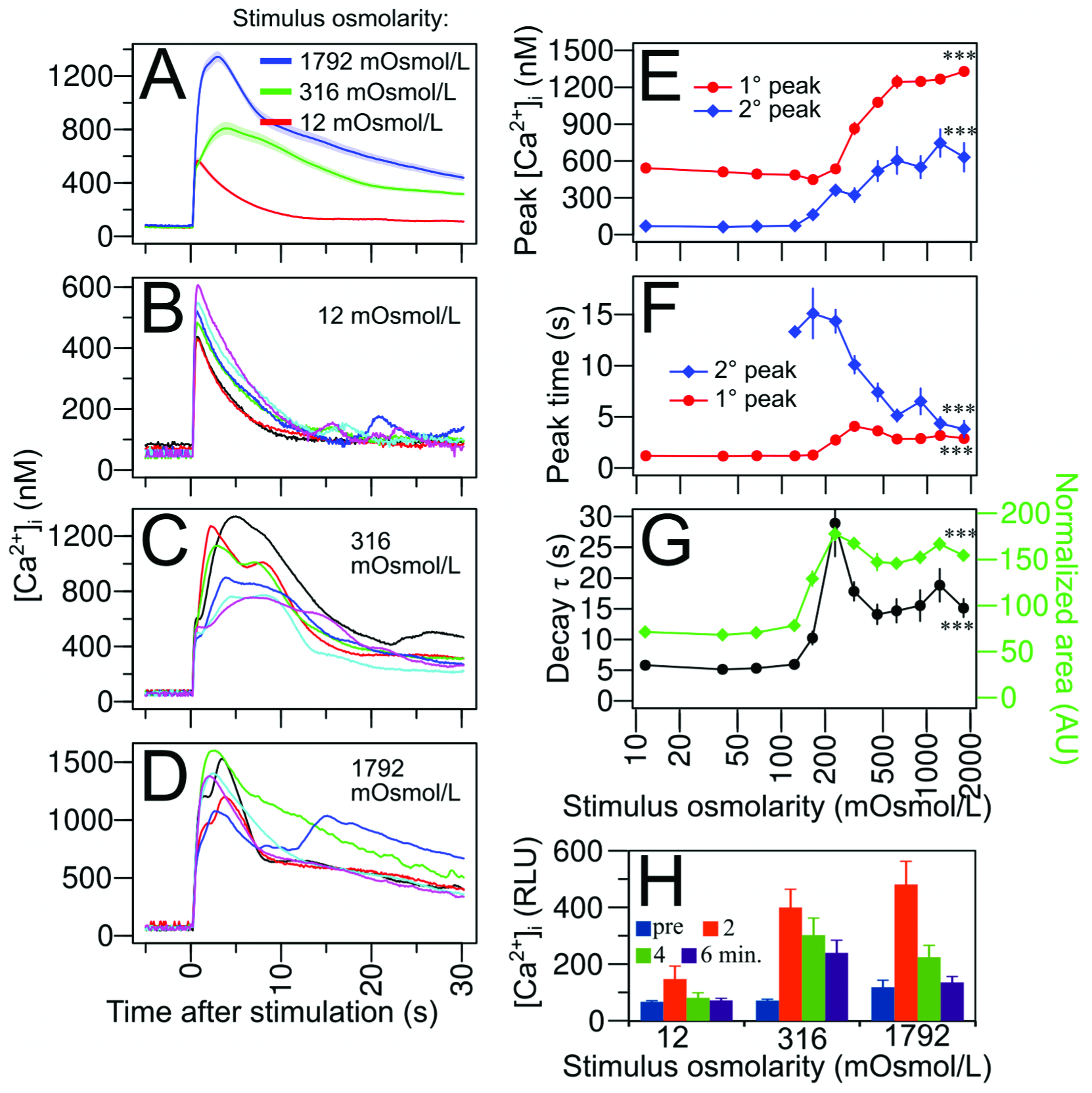
Osmotic dose-dependency of rapid Ca^2+^ response parameters. (**A**) Average [Ca^2+^]_i_ responses (solid line) ± SEM (transparent shading) of *n* = 39-40 seedlings stimulated with an injection of NaCl solutions resulting in an osmolarity of 12 (red), 316 (green), and 1792 (blue) mOsmol/L at Time = 0. (**B-D**) Six representative individual seedling Ca^2+^ responses to selected solution osmolarity. Note that the differing y-axis scales in B-D. (**E-G**) Influence of stimulus osmolarity on measured parameters of the Ca^2+^ response, including (**E**) primary (“1°”) and secondary (“2°”) peak amplitudes, (**F**) time from stimulation to the peaks, and (**G**) response termination kinetics as quantified by the decay constant “τ” (black trace) and the area under the normalized response curve (green trace). *n* = 31-40 seedlings per data point in E-G. Statistical comparisons were made between the lowest and highest stimulus concentrations using one-way ANOVA with Tukey-HSD post-hoc test. *** p<0.001. (**H**) Baseline Ca^2+^ concentrations immediately preceding (“pre”) and 2, 4, and 6 minutes following stimulation with NaCl solutions to result in the indicated osmolarity. Error bars represent ± SEM.

To determine how a dynamic osmotic environment affects hyperosmotic-induced Ca^2+^ responses, we first established how different response parameters varied as a function of stimulus osmolarity. As the log(osmolarity) of the stimulus increased, primary peaks rose in a quasi-sigmoidal fashion (Fig. 1E). The EC50, determined by the osmolarity needed to elicit 50% of the maximal response was ~316 mOsmol/L for primary (1°) peak amplitudes. The time from stimulation to primary peak increased slightly but significantly (Fig. 1F); the time from stimulation to the largest secondary peaks was reduced dramatically at higher osmolarities (Fig. 1F). Termination kinetics, measured in terms of an exponential decay constant (τ) and integrated area under normalized curves showed that termination rates slowed as the hyperosmotic stimulus was increased, but then fell slightly at very high stimulus intensities (Fig. 1G). The sensory threshold, determined by the osmolarity that achieves above-background responses, was determined to be in the range of 123-164 mOsmol/L for the parameters measured here (Figs. 1E-G).

Under the stimulation paradigm used in this study, the hyperosmotic stress is applied to a well containing the seedling, so the stress remains constant after application of the stress. However, the intracellular Ca^2+^ levels did not remain at peak elevations. Instead, the Ca^2+^ levels decayed dramatically within 30 seconds after stimulation (Figs. 1A-D), and approached prestimulus levels several minutes after stimulation for low (12 mOsmol/L) and high (1792 mOsmol/L) stimulus intensities (Fig. 1H). The intermediate (316 mOsmol/L) stimulus intensity showed a slower decay timecourse (Fig. 1G,H).

### Potentiation of rapid hyperosmotic stress-induced Ca^2+^ responses by prior hyperosmotic exposure

We next pre-exposed *Arabidopsis* seedlings to intermediate levels of hyperosmotic stress for prolonged durations (~1-3 hrs) and measured rapid Ca^2+^ responses to subsequent hyperosmotic stimuli. Interestingly, we found that raising the pre-stimulus osmolarity from 21 mOsmol/L to 149 mOsmol/L resulted in larger, more robust hyperosmotic-induced Ca^2+^ responses (Fig 2A). By quantifying the features of hyperosmotic responses as a function of starting osmolarity, we found that the primary peak amplitude was amplified by starting osmolarity, but not the secondary peak (Fig. 2B, Fig. S2C-D). Given that increasing background stress resulted in larger response amplitudes, we term this phenomenon “osmo-sensory potentiation”, after analogous potentiation phenomena described in Neuroscience (26) and Pharmacology (27). When the background stress was raised above ~212 mOsmol/L, the primary peak amplitudes were no longer amplified, but instead returned to non-potentiated levels (Fig. 2B). Sensory potentiation of hyperosmotic-induced Ca^2+^ responses consistently occurred regardless of whether the osmolyte was sorbitol or NaCl in either the pre-exposure media or the stimulation solution (Fis. S2A-B).

**Figure 2.**
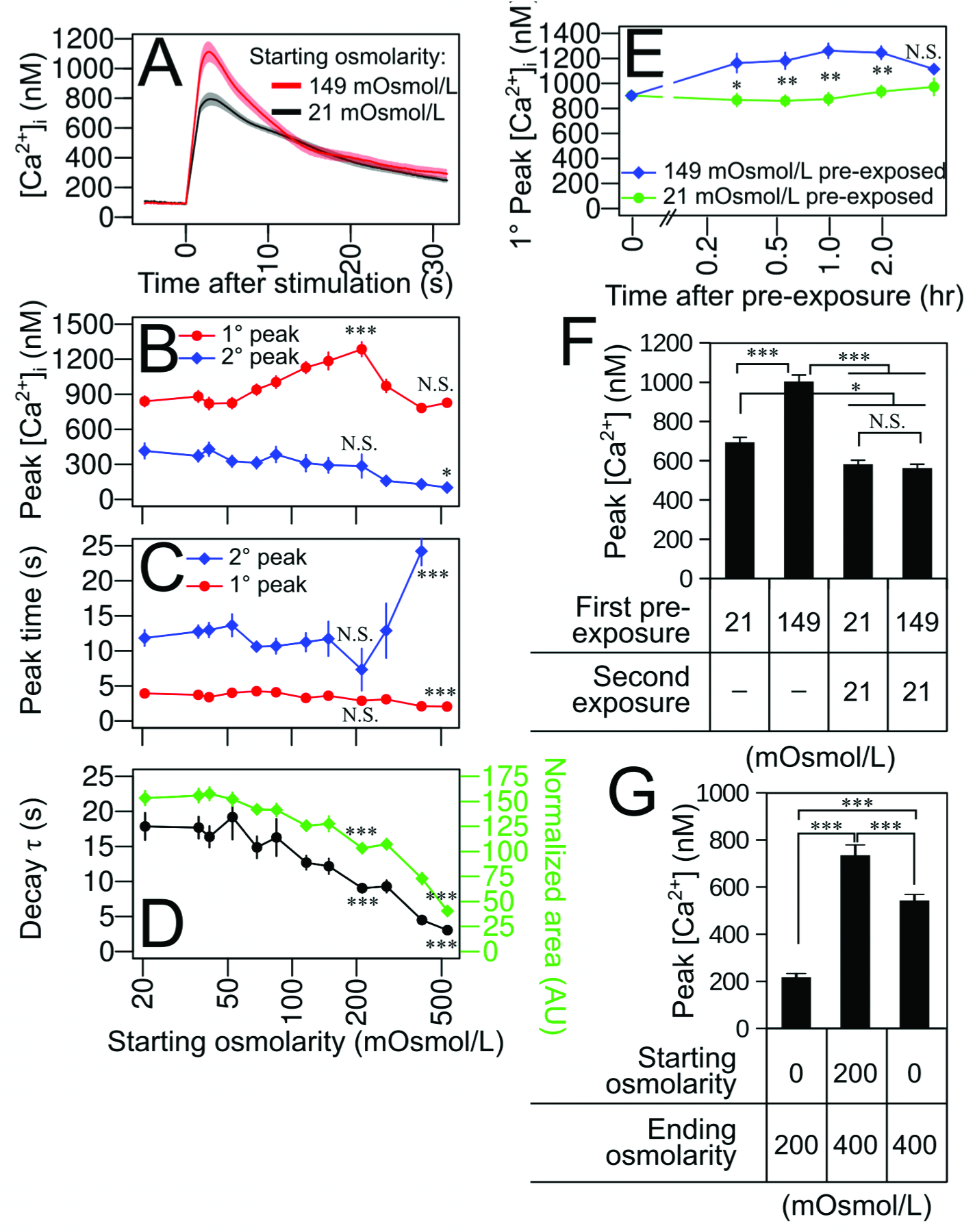
Potentiation of rapid hyperosmotic-induced Ca^2+^ responses by prior exposure of seedlings to hyperosmotic stress. (**A**) Average seedling Ca^2+^ responses (solid line) ± SEM (transparent shading) pre-exposed for 1-2 hrs to low (21 mOsmol/L, black trace), and high (149 mOsmol/L, red trace) osmolarity media, stimulated with an injection of 0.7 volumes of 700 mM sorbitol solution at Time = 0. *n* = 40 seedlings per trace. (**B-D**) Influence of starting osmolarity prior to stimulation on measured parameters of the Ca^2+^ response, including (**B**) primary and secondary peak amplitudes, (**C**) time from stimulation to the peaks, and (**D**) response termination kinetics as quantified by the decay constant “τ” (black trace) and the area under the normalized response curve (green trace). *n* = 40 seedlings per data point in (**B-D**). Statistical comparisons were made between an indicated data point and the the lowest starting osmolarity data point using one-way ANOVA with Tukey-HSD post-hoc test. (**E**) Time course of sensory potentiation, comparing primary peak amplitudes of seedlings stimulated with an injection of hyperosmotic media and pre-exposed to low (21 mOsmol/L, green trace), and high (149 mOsmol/L, blue trace) osmolarity media as a function of time from pre-exposure to time of stimulation. *n* = 62-69 seedlings per data point. Statistical comparisons were made between 21‐ and 149-mOsmol/L preexposure at each time point using one-way ANOVA with Tukey-HSD post-hoc test. (**F**) Reversibility of sensory potentiation assessed by serial exposure to either low (21 mOsmol/L) or high (149 mOsmol/L) osmolarity media. *n* = 91-93 seedlings per data point. (**G**) Ca^2+^ responses were recorded in the presence of 0 or 200 mOsmol/L starting and 200 or 400 mOsmol/L ending osmolarities imposed by sorbitol stress. *n* = 46 – 58 seedlings per condition. Error bars represent ± SEM. Statistical comparisons were made between conditions using one-way ANOVA with Tukey-HSD post-hoc test. *** p<0.001, ** p<0.01, * p<0.05, N.S p>0.05.

Pre-exposure to higher levels of hyperosmotic stress (512 mOsmol/L) resulted in a modest shortening of time to reach the primary Ca^2+^ peak (Fig. 2C). In contrast, the time to reach the secondary peak was lengthened significantly at high starting osmolarities (Fig. 2C). Pre-exposure to higher levels of hyperosmotic stress also resulted in more rapid termination kinetics and area under the normalized peaks (Fig. 2D).

Time-course analysis revealed that sensory potentiation began to occur within 18 minutes after pre-exposure to intermediate osmolarity, the sensory potentiation peaked by 2 hours after pre-exposure (Fig. 2E). To determine whether this sensory potentiation was reversible, we reexposed the seedlings a second time to low osmolarity media prior to high-osmolarity stimulation and Ca^2+^ measurement. We found the potentiation effects of the first pre-exposure to high osmolarity could be completely erased by subsequent exposure to low osmolarity media prior to hyperosmotic stimulation (Fig. 2F).

To further test how potentiation is affected by both relative and final osmolarity changes, we varied the starting (pre-treatment) and ending (stimulus) osmolarities and measured the resulting Ca^2+^ responses. We found that the seedlings starting from 200 and ending at 400 mOsmol/L showed a larger Ca^2+^ response than the 0 to 200 and 0 to 400 mOsmol/L Ca^2+^ responses (Fig. 2G). These results demonstrate that elevating the starting osmolarity potentiates the subsequent Ca^2+^ in response to the same ending osmolarity.

### Osmosensing and hyperosmotic-induced sensory potentiation occur in roots

There is some debate over the site of initial generation of rapid hyperosmotic-induced Ca^2+^ responses, which may depend on experimental conditions. While some studies show that the Ca^2+^ signal first appears in roots (9, 12), another indicates that the Ca^2+^ signal may occur simultaneously and more strongly in the shoots (15). While the Ca^2+^ response occurs in many cell types, it is reported to be largest and occurs earliest within root epidermis cells (24, 28). We measured hyperosmotic-induced Ca^2+^ responses in isolated roots, isolated shoots, and whole seedlings (Fig. 3A-B). Isolated roots displayed a larger hyperosmotic-induced Ca^2+^ response compared to shoots when exposed to the same stimulus (Fig. 3B). By varying the starting osmolarity, we determined that whole seedlings and isolated roots were competent to exhibit potentiation at a 90% confidence level, while isolated shoots were not (Fig. 3B).

**Figure 3.**
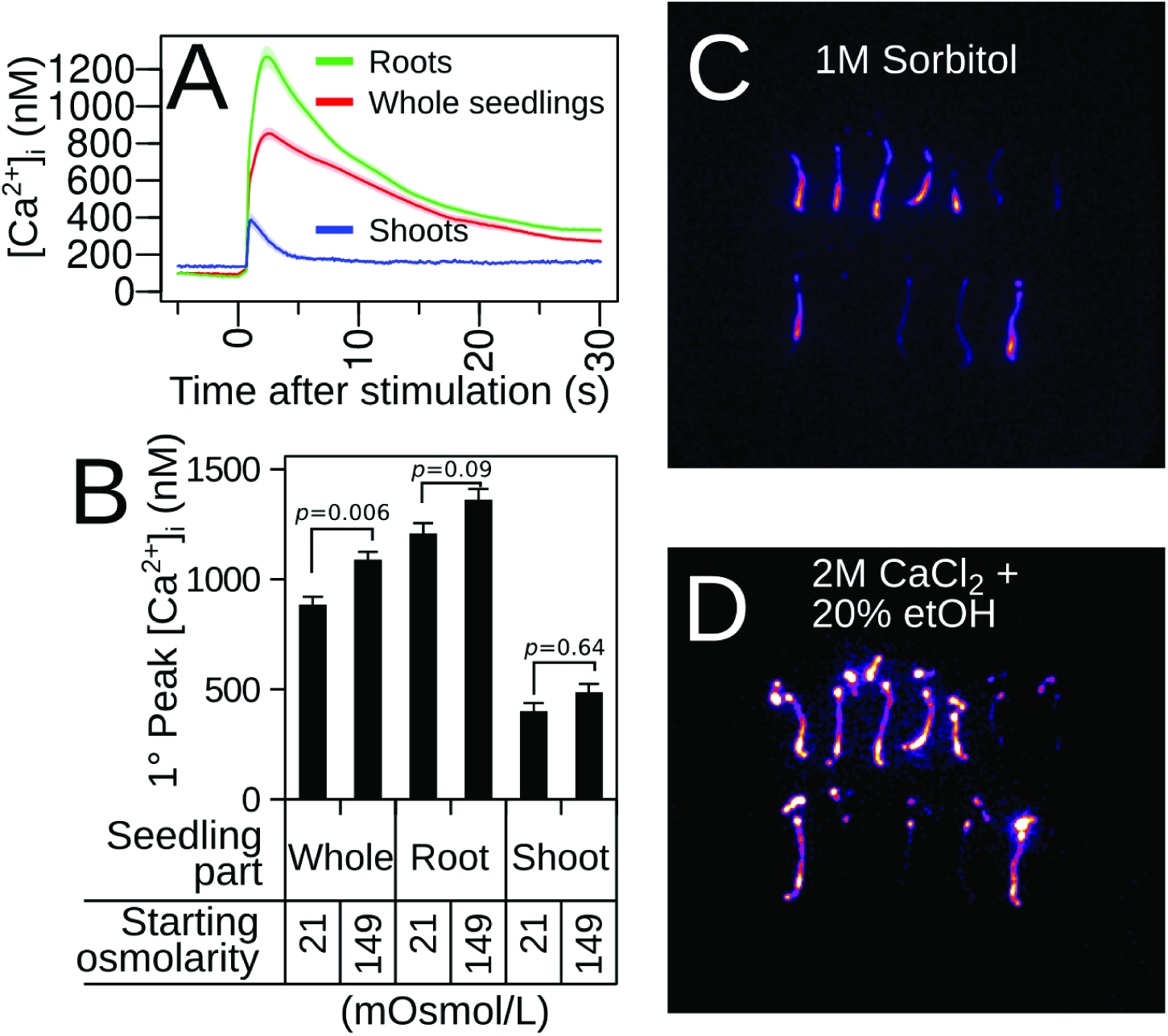
Rapid hyperosmoticinduced Ca^2+^ responses and potentiation occur primarily in roots. (**A**) Average Ca^2+^ responses (solid lines) ± SEM (transparent shading) of 1-week-old whole seedlings (red trace), isolated roots (green trace), and isolated shoots (blue trace) pre-exposed for 1-2 hrs to low (21 mOsmol/L) osmolarity media and stimulated with an injection of hyperosmotic sorbitol solution at Time = 0. *n* = 37-40 seedlings per trace. (**B**) Primary peak amplitudes whole seedlings, isolated roots, and isolated shoots stimulated with an injection of hyperosmotic media and pre-exposed to low (21 mOsmol/L, green trace), or moderate (149 mOsmol/L, blue trace) osmolarity media. Error bars represent ± SEM. Statistical comparisons were made between starting osmolarity conditions using one-way ANOVA with Tukey-HSD post-hoc test. P-values are indicated. (**C-D**) Pseudo-colored photos depicting light emission of intact whole seedlings in response to (**C**) hyperosmotic shock (1 M sorbitol, signal integrated for 30 seconds) and subsequently to (**D**) 2M CaCl_2_ + 20% ethanol, which discharges total remaining aequorin, signal integrated for one minute.

To more directly analyze the spatial distribution of Ca^2+^ dynamics in whole, intact seedlings, plants were stimulated uniformly with hyperosmotic stress and imaged with a CCD camera (Fig. 3C). Luminescence from the aequorin reporter emitted almost exclusively from the roots, most strongly in the zones close to the root tip (Fig. 3C). A lack of light emission from the leaves could not be explained by a lack of aequorin reporter in the leaves, since subsequent discharge of total remaining aequorin with 2M CaCl_2_ + 20% ethanol revealed an abundance of active aequorin reporter remaining in leaves (Fig. 3D).

### Additive effects of abscisic acid signaling and hyperosmotic-induced potentiation

One possible mechanism for sensory potentiation could involve positive feedback of downstream hyperosmotic-induced signaling components on the upstream osmosensory components. Given that osmotic stress induces abscisic acid (ABA) biosynthesis (29, 30), and ABA enacts a multitude of physiological changes including modulation of ion channel activity and transcriptional reprogramming (31), we tested the effects of exogenous ABA application on hyperosmotic-induced Ca^2+^ responses. In 6 of 12 experiments, pre-exposure of seedlings to ABA resulted in larger-amplitude hyperosmotic-induced Ca^2+^ responses (Fig. 4A-C). In the other 6 of 12 experiments, no effect of adding exogenous ABA was observed. In experiments where ABA resulted in larger-amplitude hyperosmotic-induced Ca^2+^ responses, 2 μM ABA was sufficient to reach saturation, as 10 μM ABA did not further increase the response amplitudes (Fig. 4A-B). To determine whether this ABA effect may interact with the potentiation of the response by elevated starting osmolarity, we pre-exposed seedlings to either low or moderate hyper-osmolarity media which either lacked or contained ABA. We found that the amplitude increases induced by either ABA pre-exposure or hyperosmotic pre-exposure alone were additive when the seedlings were pre-exposed to both ABA and high starting osmolarity media (Fig. 4C); two-way ANOVA (32) analysis indicated that ABA pre-exposure together with starting osmolarity pre-exposure significantly increased the primary peak amplitudes (*p* = 2.3x10^-4^ and p= 6.79x10^-8^, respectively), but there was no detectable interaction between the ABA and starting osmolarity pre-treatments (p= 0.89). This result suggests that potentiation of the hyperosmotic-induced Ca^2+^ response by pre-exposure to hyperosmotic stress occurs by mechanisms that are at least partially distinct from those involved in ABA signaling.

**Figure 4.**
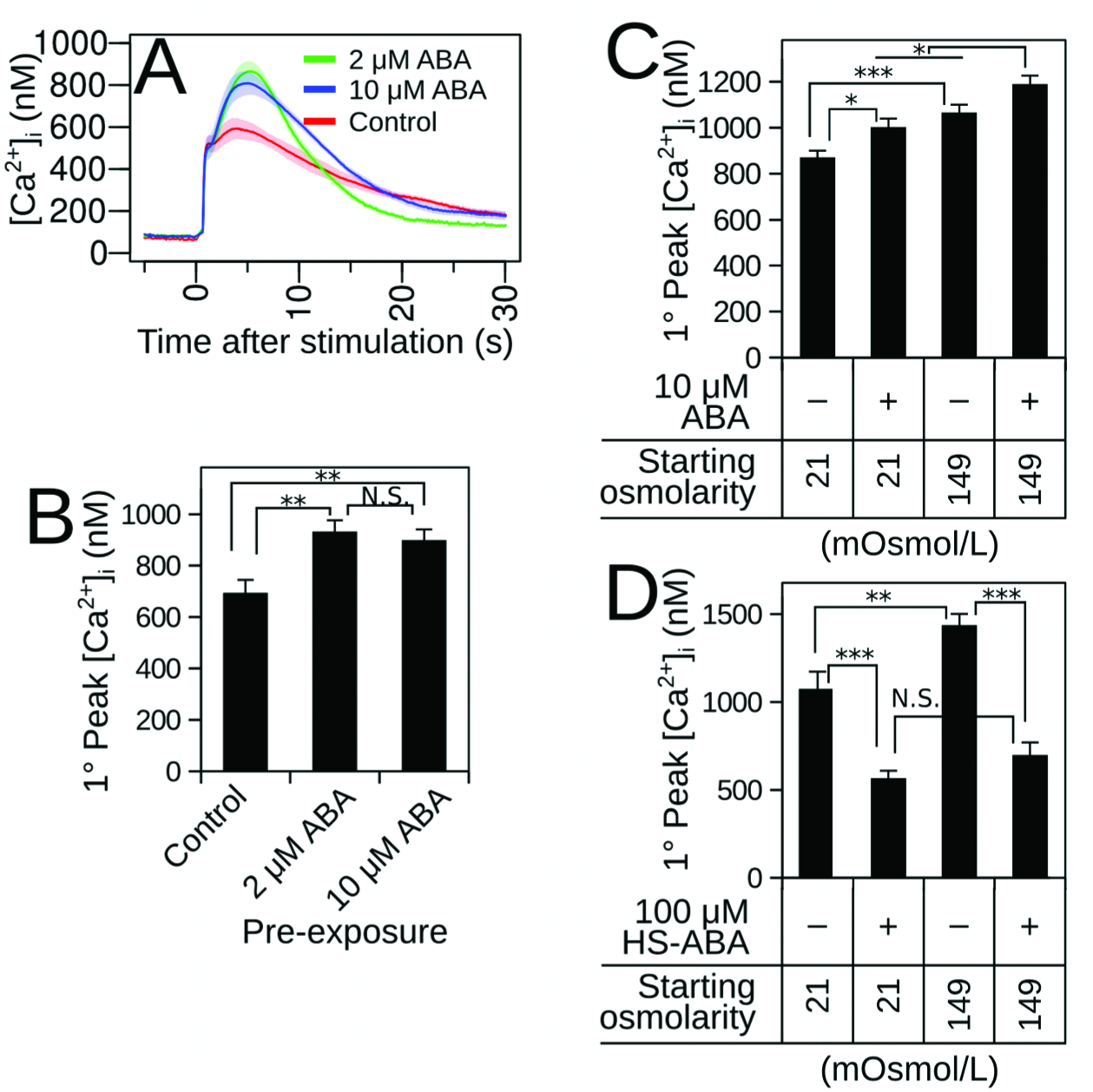
Influence of Abscisic Acid (ABA) on Ca^2+^ response amplitudes and potentiation. (**A**) Average Ca^2+^ responses (solid line) ± SEM (transparent shading) of 1 week-old whole seedlings pre-exposed for 1-2 hrs to low (21 mOsmol/L) osmolarity media with or without ABA at indicated concentrations and stimulated with an injection of hyperosmotic sorbitol solution at Time = 0. *n* = 20-23 seedlings per trace. (**B**) Quantification of primary peak amplitudes in (**A**). Statistical comparisons were made between conditions using one-way ANOVA with Tukey-HSD post-hoc test. ** p<0.01, N.S p>0.05. (**C**) Additive effects of pre-exposure to ABA and elevated osmolarity on Ca^2+^ responses in seedlings stimulated with an injection of hyperosmotic media. *n* = 47-48 seedlings per data point. Two-way ANOVA analysis revealed that ABA pre-exposure and starting osmolarity pre-exposure significantly increased the primary peak amplitudes (*p* = 2.3x10^-4^ and p= 6.79x10^-8^, respectively), but there was no detectable interaction between the treatments (p= 0.89). Depicted p-values were calculated by the Tukey-HSD post-hoc test. *** p<0.001, * p<0.05. (**D**) Effects of the ABA antagonist HS-ABA (100 μM) on hyperosmotic-induced Ca^2+^ response amplitudes in seedlings pre-exposed to low or high osmolarity media. *n* = 30-32 seedlings per data point. Error bars represent ± SEM. Two-way ANOVA revealed that starting osmolarity and HS-ABA treatment had large effects on primary peak amplitudes (*p* = 1.8x10^-4^ and *p* < 2x10^-16^, respectively). No significant interaction was seen between HS-ABA treatment and starting osmolarity. Depicted p-values were calculated by the Tukey-HSD post-hoc test. *** p<0.001, ** p<0.01, * p<0.05. The influence of ABA on the Ca^2+^ response amplitudes was observed in 6 of 12 experiments. Panels A and B are one experiment, and Panel C is another experiment.

To address this question further, we made use of a recently-developed ABA signaling antagonist, hexasulfanyl-ABA (HS-ABA). This ABA analog has been shown to bind the PYR/PYL family of ABA receptors and disrupt the receptor interactions with inhibitory PP2C proteins, thereby leading to constitutive repression of ABA signaling (33). HS-ABA drastically reduced the amplitudes of the hyperosmotic-induced Ca^2+^ responses, eliminating any discernible effect of hyperosmotic pre-treatment potentiation (Fig. 4D). Two-way ANOVA demonstrated that starting osmolarity and HS-ABA treatment had large effects on primary peak amplitudes (*p* = 1.8x10^-4^ and *p* < 2x10^-16^, respectively). Nevertheless, no significant interaction was found between HS-ABA treatment and starting osmolarity (i.e., the treatments appeared additive) (*p* = 0.07). If the effects of HS-ABA are indeed resulting from specific inhibition of ABA signaling, this result could indicate that a certain level of basal ABA signaling is a prerequisite for maintaining responsiveness of the osmosensory components regardless of potentiation status.

### Survey of ion transporters that may play a role in elicitation of hyperosmotic-induced Ca^2+^ responses

Only one mutant, OSCA1, has been described to date that impairs the rapid osmotic stress-induced Ca^2+^ elevation (15). Previous studies suggest that the Ca^2+^ stores responsible for rapid hyperosmotic-induced Ca^2+^ influx may include both the apoplast as well as intracellular organelles, since extracellular Ca^2+^-signaling inhibitors La^3+^, Gd^3+^, and EGTA as well as several intracellular Ca^2+^-signaling inhibitors all affect the amplitudes and kinetics of the osmotic-induced Ca^2+^ response (5, 9). We tested a variety of pharmacological agents and found that LaCl3, GdCl3, Amiloride, and D-cis-Diltiazem dampened the rapid osmotic-induced Ca^2+^ response (Fig. S3, Table S1), but DNQX, CNQX, D-AP5, Verapamil, Tetracaine, Ruthenium Red, U-73122, H-8, Methylene Blue, NS-2028, and ODQ did not dampen the rapid osmotic-induced Ca^2+^ response (Table S1).

We assessed candidate mutant lines for altered hyperosmotic-induced Ca^2+^ responses, including mechano-sensitive plasma membrane-localized channels. No significant effects were observed in the higher-order mechanosensitive Ca^2+^ channel root *Mid1-complementing activity mca1;mca2* (AT4G35920, AT2G17780) double mutant (34, 35) or the plasma membrane-targeted members of the *MscS-like* family *msl4;5;6;9;10* (AT1G53470, AT3G14810, AT1G78610, AT5G19520, AT5G12080) quintuple mutant (36) (Fig. S4). Additionally, it has been previously shown that the *tpc1* (AT4G03560) mutants in the vacuolar Ca^2+^-activated Ca^2+^-permeable SV channel do not affect the rapid osmotically-induced Ca^2+^ rise, but rather impaired the ensuing root-to-shoot Ca^2+^ wave (12).

It is recently becoming evident that plastids represent a significant pool for Ca^2+^ release in plants (21, 37, 38); we measured hyperosmotic-induced Ca^2+^ responses in plastidial-localized potassium/proton antiporter *kea* mutant lines. KEA1 (AT1G01790) and KEA2 (AT4G00630) are targeted to the inner plastid envelope membrane and are required for plastid ion homeostasis and osmo-regulation (39, 40). Loss of function results in morphologically swollen plastids and decreased photosynthetic activity (41). Since KEA1 and KEA2 act as K+/H+ antiporters, mutation of these genes is expected to perturb the electrochemical potential of these—and possibly other—ions across the envelope membrane. We found that *kea1-2kea2-2* double mutants displayed a substantially reduced amplitude of the rapid hyperosmotic-induced Ca^2+^ response, especially at high starting osmolarity (Fig. 5A-B). Comparing conditions of low and high starting osmolarity, we found that the *kea1-2kea2-2* double mutant line was competent to exhibit sensory potentiation, despite overall reduced amplitudes compared to wild-type (Fig 5A-B). Transgenic expression of wild-type *KEA2* under the control of the AtUBQ10 promoter was sufficient to rescue the low amplitude Ca^2+^ phenotype of the *kea1-2kea2-2* double mutant (Fig. 5E-F).

**Figure 5.**
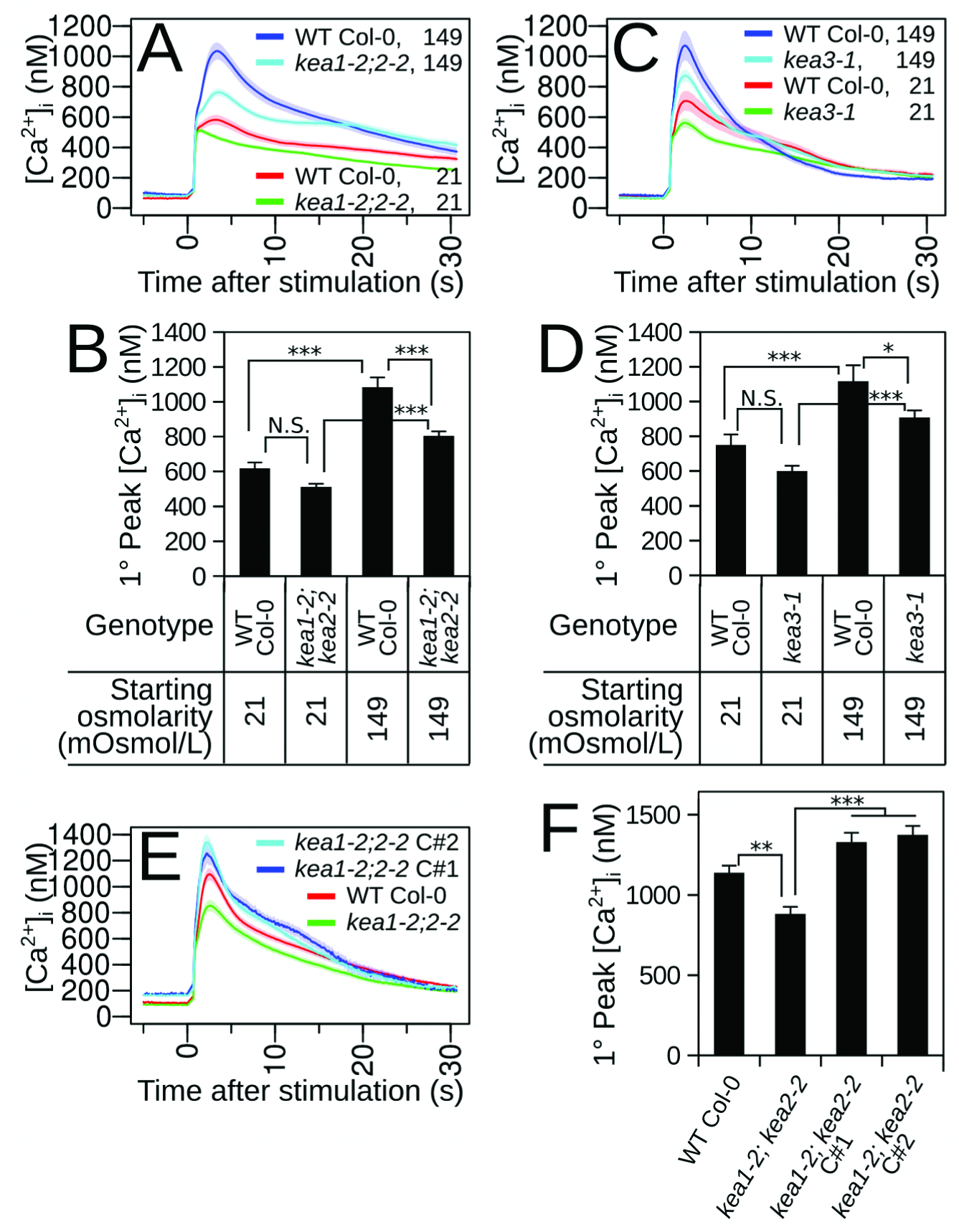
Plastidial *kea* mutations reduce rapid hyperosmotic-induced Ca^2+^ response amplitudes, but do not eliminate sensory potentiation. (**A**) Average Ca^2+^ responses (solid line) ± SEM (transparent shading) of 1-week-old wild-type (Col-0) or *kea1-2; keci2-2* double mutant seedlings preexposed for 1-2 hrs to low (21 mOsmol/L) or high (149 mOsmol/L) osmolarity media and stimulated with an injection of hyperosmotic sorbitol solution at Time = 0. *n* = 44-64 seedlings per trace. (**B**) Quantification of primary peak amplitudes in (**A**). Two-way ANOVA demonstrated that the effects of both Genotype and Starting Osmolarity on primary peak amplitudes were highly significant (*p* = 2.77l0^-7^ and 2.0l0^17^, respectively), and the interaction between genotype and starting osmolarity was also significant (*p* = 0.01). Statistical comparisons were made with Tukey-HSD post-hoc test.(**C**) Average Ca^2+^ responses (solid line) ± SEM (transparent shading) of 1-week-old wild-type (Col-0) or *kea3-1* mutant seedlings preexposed for 1-2 hrs to low (21 mOsmol/L) or high (149 mOsmol/L) osmolarity media and stimulated with an injection of hyperosmotic sorbitol solution at Time = 0. *n* = 28-66 seedlings per trace. (**D**) Quantification of primary peak amplitudes in (**C**). Two-way ANOVA demonstrated that the effects of both Genotype and Starting Osmolarity on primary peak amplitudes were significant (*p* = 4.5x10^-4^ and 6.69x10^-10^, respectively), but the interaction between Genotype and Starting Osmolarity was not significant (*p* = 0.59). Statistical comparisons were made with Tukey-HSD post-hoc test. (**E-F**) Genetic complementation with a wild-type copy of *Kea2* can rescue the reduced amplitude phenotype of *kea1-2;kea2-2* double mutant. Error bars represent ± SEM. Statistical comparisons were made using one-way ANOVA with Tukey-HSD post-hoc test. *** p<0.001, ** p<0.01, * p<0.05, N.S p>0.05.

We next tested a mutant in the *KEA3* (AT4G04850) gene; KEA3 is localized to the thylakoid membrane (41, 42)—the actual Ca^2+^ storage site within the plastid. *Kea3* mutants show morphologically intact chloroplasts, but mutation results in altered photosynthetic energy absorption under dynamic light conditions (42). Interestingly, we found that the *kea3-1* mutant line displayed a reduced amplitude of the hyperosmotic-induced Ca^2+^ response, much like the *kea1-2kea2-2* double mutant line (Fig. 5C-D). Likewise, the *kea3-1* mutant line was still competent to exhibit sensory potentiation (Fig. 5C-D). Thus the impairment in osmotic-induced Ca^2+^ cannot be solely linked to aberrant chloroplast morphology of the *kea1kea2* mutant, but correlates with effects of both mutants on chloroplast ion homeostasis and photosynthetic efficiency.

## Discussion

Here we report that early sensory components leading to rapid osmotic-induced Ca^2+^ responses in plants can be up-regulated (“potentiated”) by prior exposure to osmotic stress, while mutation of plastidial KEA transporters results in a reduced rapid Ca^2+^ elevation. Given that plants exhibit rapid hyperosmotic-induced Ca^2+^ responses with or without pre-exposure to osmotic stress, this indicates that the sensory components is always present and functional. However, the expression, activity, or sensitivity of the sensory components is increased after experiencing previous hyperosmotic stress, thereby exhibiting sensory potentiation. Important and interesting examples exist in the literature describing enhancement of responses after experiencing stress, which is sometimes referred to as “priming” or “memory” (43–46). The present study, however, is the first to report that the earliest rapid osmotic stress response sensory components can be up-regulated in such a manner. Sensory potentiation, as described here, represents a distinct case from these previously described examples since all responses downstream of the initial perception event could be affected. Moreover, the rapid onset and reversibility of sensory potentiation suggest that this phenomenon may be of key importance for “on-the-fly” adjustment of the sensitivity to osmotic conditions as experienced by plant roots in soils when experiencing osmotic stress.

We found that under constant hyper-osmotic stimulation, the Ca^2+^ response returns to baseline. This return to baseline suggests that negative feedback mechanisms must exist to inactivate sensory components, including closing Ca^2+^ channels and removal of cytosolic Ca^2+^ by active transport (47). Many sensory systems exhibit a similar return to baseline, which is termed “sensory adaptation” (48); examples include mammalian vision and audition, bacterial chemotaxis, and yeast osmoregulation. These require integral feedback control to balance sensory components' activation with inactivation (49, 50). Returning to baseline ensures that the system remains responsive to further relative changes in stimulus intensity.

While we had initially hypothesized that ABA elevation induced by osmotic stress could feed back to up-regulate the osmo-sensory components, we found that the sensitivity increase caused by ABA is more variable and largely independent of the sensitivity increase caused by prior exposure to hyperosmotic stress (Fig. 4). Despite our best efforts to control environmental conditions, the effects of exogenous ABA application varied experiment-to-experiment nonetheless. Given that ABA only had an effect in half of our tested experiments and given that HS-ABA could reduce the amplitude of osmotic-induced Ca^2+^ responses, we hypothesize that a basal amount of ABA is required for osmotic-induced Ca^2+^ responses, and once that basal level is reached, no more ABA-dependent potentiation can occur. This hypothesis can be tested further in the future with mutant lines that are deficient in ABA biosynthesis. We observed a rapid (< 20-minute) onset of potentiation by pre-exposure to hyperosmotic stress (Fig. 2E), but it has been observed that osmotic-induced ABA increases in roots occur on a substantially slower timescale (51), consistent with other findings. Therefore, it is unlikely that a rapid increase in ABA concentration can account for osmotic-induced potentiation.

Given that hyper-osmotic stress results in an increase in [Ca^2+^]_i_, it raises the question of which Ca^2+^ stores are important for this response. Our analyses of the *mca1;mca2* double mutant and the *msl* quintuple mutant show that other channels are required to display the rapid wild-type osmotic-induced Ca^2+^ responses. Our analyses of the plastidial KEA transporters offer initial insight into this question. KEA1 and KEA2 are localized to the inner envelope of the plastid, whereas KEA3 is localized to the thylakoid membrane (39, 41, 42). While *kea1-2kea2-2* double mutants display phenotypes associated with severely impaired chloroplast functions (41), the *kea3-1* mutant line does not affect chloroplast morphology and is only known to display electron transport phenotypes under dynamic light situations (42). In this study, we found that both *kea1-2kea2-2* double mutants and *kea3-1* mutants showed reduced osmotic-induced Ca^2+^ responses despite being grown in standard lighting conditions. These results offer the tantalizing prospect that plastidial membrane energetics and Ca^2+^ stores may be linked and required for wild-type hyperosmotic-induced Ca^2+^ responses (21, 52). In animal systems, for instance it has been demonstrated that mitochondria play an important function in Ca^2+^ release (53, 54). A recent study shows that *KEA1*, *KEA2*, and *KEA3* transcripts are detectable in roots (55). Additionally, root leukoplasts have intracellular membranous lamellar vesicles and plastoglobules (56), the latter of which has significant association with thylakoids in terms of proteome composition (57). Thus it is possible that the calcium signaling phenotype found in the KEA mutants is due to disrupted root plastid ion homeostasis, but presently an indirect effect (leaf-to-root) or systemic effect cannot be excluded, which will require further analyses. The effects of these mutations could be more indirect, affecting ion homeostasis or osmotic perception at the cell surface or in other organelles. Note that mutations in the kea3 transporter do not visually affect chloroplast ultra-structures (41, 42). It has been reported that disrupted plastid osmoregulation in the *msl2msl3* double mutant activates drought-related phenotypes in the absence of an external stress (58). The present study highlights that the involvement of plant plastids in the initial osmotic sensory events deserves further attention.

A novel family of osmotically-gated Ca^2+^ ‐permeable ion channels has been reported in plants (15, 18), and mutations of one family member are the only mutations known to reduce hyperosmotic-induced Ca^2+^ responses primarily in leaves (15). Our study identifies the regulation of the osmosensory components by preceding exposure to hyperosmotic stress (potentiation), by the hormone ABA, and by the activities of plastid-localized KEA transporters. These results suggest that the initial osmotic perception events leading to induction of rapid [Ca^2+^]_i_ elevations represent a complex trait that is influenced by several factors. Together, this work will guide mechanistic studies toward elucidating the osmo-sensory components in plants.

## Materials and Methods

### *Arabidopsis* plant lines

Wild-type Col-0 *Arabidopsis* seedlings harboring a T-DNA insertion carrying a 35S:Apoaequorin expression cassette were used throughout this study. The lines were either direct descendants of the original kanamycin-resistant “pMAQ2” line (59) (shared by M. Knight, Durham University, and Z.M. Pei, Duke University) or an independently-generated hygromicin-resistant line (shared by A. Dodd, Bristol University). Both lines showed nearly identical expression levels of Apoaequorin, and sensory potentiation was confirmed to occur in both lines. The hygromicin-resistant 35:Apoaequorin line was crossed to the *kea1-2;2-2* (SAIL_1156_H07;

SALK_020285) double mutant and the *kea3-1* (SAIL_556_E12) single mutant lines (41). The resulting F1 individuals were self-pollinated, and F2 progeny were genotyped and assayed for aequorin expression (41). The F2 individuals that were homozygous for the intended *kea* mutations as well as expressing aequorin were propagated to the F3 generation and tested for hyperosmotic-induced Ca^2+^ responses. Four F3 lines were tested with similar results. *Mca1;2* mutants, provided by Hidetoshi Iida, and *Msl4;5;6;9;10* mutants, provided by Elizabeth Haswell, were transformed by the agrobacterium-mediated floral dip method using a “pMAQ2-NEW” vector provided by Zhen-Ming Pei. T1 transformants were selected based on BASTA resistance and aequorin expression. Two T2 lines were tested for each mutant combination. For the *Kea* complementation experiment, the *kea1-2;2-2* double mutant expressing aequorin was transformed by the agrobacterium-mediated floral dip method using a vector expressing Kea2-mVenus fusion protein under the control of the UB10 promoter (41). T1 transformants were selected based on enhanced hyg resistance as well as non-chlorotic leaves, indicating functional complementation. Two T2 lines were tested for osmotic-induced Ca^2+^ responses, analyzing the ˜¾ green seedlings as segregating complemented individuals.

### Luminometry measurements of Ca^2+^ responses in seedlings

Ethanol-sterilized seeds were plated individually in wells of sterile, white, 96-well microplates containing 130 μl media composed of 1/2 Murashige and Skoog (MS) Media (2.2 g/L) + 0.5 g/L 2-(*N*-morpholino)ethanesulfonic acid (MES) + 1% sucrose + 0.08% Phyto Agar. The low percentage of Phyto Agar provided modest support for seedling growth while allowing subsequent rapid mixing of injected pre-treatment solutions and stimulus solutions. The clear lids were sealed with Parafilm to prevent media evaporation. The seedlings were cold stratified for 2-3 days at 4°C in the dark, and were then grown for 6.5 days in a growth cabinet (16h light, 8h dark, 22°C, 60-80 μEm^-2^s-^1^ light intensity). Twelve to 14 hours prior to the experiment, 130 μL of water containing 2 μM native coelenterazine (Ctz) (Promega) was added to each well. The lid was placed back on the plate, but was not sealed with Parafilm again. The plate was covered by aluminum foil to prevent light-induced degradation of Ctz, and was incubated overnight in the growth cabinet.

The following day, just prior to experiments, 130 μL of media was removed from each well, and 130 μL of pre-exposure media was added. Unless otherwise noted, the pre-exposure proceeded for ~1 hr prior to stimulation, and given that the luminometer recordings take ~45 minutes to measure all of the wells, the pre-exposure generally lasted for 1-1.75 hrs. In the case of the serial pre-exposure experiment (Fig. 2F), the first pre-exposure proceeded for 1hr, and the second pre-exposure was done immediately prior to luminometry readings. Therefore, the second pre-exposure proceeded ~5 ‐45 minutes before a given well was stimulated and recorded. In the case of the time-course measurements (Fig. 2E), the wells were pre-exposed with staggered start times, factoring in the additional time that the plate reader took to reach a given well during measurement. Unless otherwise noted, the pre-exposure media was water, resulting in a final diluted osmolarity with the starting media (41 mOsmol/L) of 21 mOsmol/L. In the case of potentiated responses, unless otherwise noted, the pre-exposure media was 256 mM sorbitol, resulting in a final diluted osmolarity with the starting media of 149 mOsmol/L. In the experiments involving ABA, HS-ABA, pharmacological agents, and various starting osmolarities (Fig. 4), the pre-exposure solutions contained 2x the final diluted concentrations of the compounds. The HS-ABA was synthesized as a racemic alkyl thioether according to (33, 60). In the experiments evaluating pharmacological inhibition of osmotic-induced Ca^2+^ responses, the compounds were dissolved in stock solvents indicated in Table S1, and then diluted in their final concentration in water and the media in the well to their final concentrations indicated in Table S1. These solutions were incubated for 30 minutes prior to stimulation. The drug concentrations used were determined from previously-reported studies as well as our own dose-response analyses. In all cases, just prior to luminometry recordings, 130 μL (0.5 volumes) of the media was removed to allow enough empty volume for stimulation solutions.

The 96-well plates containing seedlings were loaded into a Berthold Mithras LB930 luminometer plate reader. Luminescence was measured every 0.1 seconds for 5 seconds to establish a baseline reading, and then 100 μL of stimulation solution was automatically injected at the device's highest rate to ensure the shortest disruption of recording. Unless otherwise noted, the injected stimulation solution was 700 mM sorbitol. In the case of the dose-response measurements in Fig. 1, stimulation was done with NaCl due to solubility limits of sorbitol. The lower osmolarity dose-dependencies were confirmed with sorbitol stimulus. After injection, light was measured every 0.1 seconds for an additional 30 seconds to one minute. Due to mechanical limitations of the luminometer, there was generally a period of 0.2 seconds during injection when light could not be measured. The data from these recordings were exported as tab-separated values spreadsheets and were designated as the “stimulus-induced” luminescence data.

After all 96 wells had been stimulated and measured, the plate was removed from the luminometer and 100 μL of solution were removed from each well. The plate was again loaded into the luminometer. Luminescence was measured every 0.1 seconds for 2 seconds to establish a baseline reading, and then 100 μL of 2 M CaCl_2_ + 20% ethanol solution was automatically injected at the device's highest rate to ensure the shortest disruption of recording. After injection, light was measured every 0.1 seconds for an additional 15-30 seconds. The data from these recordings were exported as tab-separated values spreadsheets and were designated as the “aequorin discharge” luminescence data.

### Quantification of response features

A data analysis script was written in R (61) to quantify features of individual responses as well as averages of conditions or genotypes (Supplemental Information). Raw light emission was converted to Ca^2+^ concentration by the empirically-determined formula pCa = 0.332588(-logk) + 5.5593, where k is a rate constant equal to stimulus-induced luminescence counts per second divided by total remaining counts (elicited by discharge by 2M CaCh + 20% ethanol) (25). It should be noted that this calculation assumes that all cells containing aequorin are responsive to the stimulus. Since this assumption may not always hold true, the calculated apparent [Ca^2+^]_i_ may be an underestimate of the true [Ca^2+^]_i_. The [Ca^2+^] reported in this study reflect apparent [Ca^2+^]_i_.

Peak detection was performed using the Peaks library (62), using empirically-established values that generally fit the observed peaks. The primary (1°) peak was defined as the first observable peak subsequent to stimulation. The secondary (2°) peak was defined as the largest detected peak that was not the primary peak. The largest peaks were normalized to a maximum value of 1, and were fit to the single exponential decay function R = exp(-T/τ), where R is the response amplitude at time T, and τ is the decay time constant. Given that the termination kinetics were much slower than activation kinetics, a calculation of integrated area under the normalized peak also served as a useful metric of termination kinetics. Each of the calculated features were plotted in “beeswarm” stripcharts depicting individuals, averages, and confidence intervals (63).

### Imaging of whole-seedling Ca^2+^ responses

Ethanol-sterilized seeds were plated on media containing 1/2 MS media (2.2 g/L) + 0.5 g/L MES + 1.5% Phyto Agar. The plates were wrapped in Micropore tape and were cold stratified for 2-3 days at 4°C in the dark. The plates were transferred to a growth cabinet, and the seedlings were grown vertically for 6.5 days under 16h light, 8h dark, 22°C, 60-80 μEm^-2^s-^1^ light intensity. The evening prior to the experiment, the plate was repositioned from vertical to lie flat horizontally. A few drops of 37°C 0.5% low-melt agarose were added along the length of the seedlings (both shoots and roots) to affix the seedlings to the plate. After the agarose solidified, 20 mL of water containing 1 μM native Ctz was added to the plate. The plate was covered in aluminum foil and placed back in the growth cabinet overnight.

The following day, the solution containing Ctz was decanted off the plate, and the plate was placed horizontally in an imaging cabinet. A CCD camera was used to image the light emitting from the seedlings. For sorbitol stimulation, 25 mL of 1M sorbitol was dispensed onto the seedlings, and the photos integrated light for 30 seconds. The sorbitol stimulation solution was decanted off the the plate, and the plate was again placed horizontally in the imaging cabinet. Twenty-five mL of 2M CaCl2 + 20% ethanol was dispensed onto the seedlings, and the photos integrated light for 1-2 minutes. The photos were subsequently pseudo-colored in ImageJ with the “Fire” lookup table to emphasize light emission intensities.

## Acknowledgements

We thank Antony Dodd, Zhen-Ming Pei, and Marc Knight for Aequorin-expressing *Arabidopsis* seed lines and vectors. Jose Pruneda-Paz and Katia Bonaldi for support with imaging of seedlings. Felix Hauser provided guidance in data analysis. Elizabeth Haswell provided the *msl* quintuple mutant. Hidetoshi Iida provided the *mca1;mca2* double mutant. A.B.S. was supported by a fellowship through the Life Sciences Research Foundation funded by the U.S. Department of Energy, Office of Science, Office of Basic Energy Sciences, Physical Biosciences. This project was funded by grants from the National Institutes of Health (GM060396-ES010337) and the National Science Foundation (MCB1414339) to J.I.S. H.-H.K. was supported by a Human Frontier Science Program Long-Term fellowship and an Alexander von Humboldt Feodor Lynen fellowship.

---

**Table S1.** Effects of pharmacological inhibitors on rapid osmotic-induced Ca^2+^ responses.

| Drug | IUPAC Name | CAS | [ ], solvent | Putative targets | Effect |
| --- | --- | --- | --- | --- | --- |
| Lanthanum chloride | LaCl<sub>3</sub> | 10099-58-8 | 0.06–32mM, water | Ca<sup>2+</sup> channels | Reduced amplitude (≥ 0.125 mM) |
| Gadolinium chloride | GdCl<sub>3</sub> | 10138-52-0 | 1 mM, water | Ca<sup>2+</sup> channels | Reduced amplitude |
| Amiloride | 3,5-diamino-6-chloro-N-(diaminomethylene)pyrazine-2-carboxamide | 2016-88-8 | 0.037 – 1.3 mM, water | cyclic nucleotide-gated channels, epithelial sodium channel, Na<sup>+</sup>/H<sup>+</sup> and Na<sup>+</sup>/Ca<sup>2+</sup> exchangers, Acid-sensing ion channels, ATP-sensitive K<sup>+</sup> channels | Reduced amplitude (≥ 0.7 mM) |
| D-*cis*-Diltiazem | (2S,3S)-(+)-*cis*-3-Acetoxy-5-(2-dimethylaminoethyl)-2,3-dihydro-2-(4-methoxyphenyl)-1,5-benzothiazepin-4(5H)-one hydrochloride | 33286-22-5 | 50, 100 μM, water | cyclic nucleotide-gated channels, L- and P-type Ca<sup>2+</sup> channels, 5-HT<sub>3</sub> receptor | Reduced amplitude (100 μM) |
| DNQX | 6,7-Dinitroquinoxaline-2,3-dione | 2379-57-9 | 1 mM, DMSO | ionotropic glutamate receptors | No effect |
| CNQX | 7-nitro-2,3-dioxo-1,4-dihydroquinoxaline-6-carbonitrile | 115066-14-3 | 1 mM, DMSO | ionotropic glutamate receptors | No effect |
| D-AP5 | D-(-)-2-Amino-5-phosphonopentanoic acid | 79055-68-8 | 1 mM, water | ionotropic glutamate receptors | No effect |
| Verapamil | (RS)-2-(3,4-Dimethoxyphenyl)-5-[[2-(3,4-dimethoxyphenyl)ethyl]-(methyl)amino]-2-prop-2-ylpentanenitrile | 52-53-9 | 50 μM, water | L-type Ca<sup>2+</sup> channels | No effect |
| Tetracaine | 2-(dimethylamino)ethyl 4-(butylamino)benzoate | 94-24-6 | 100 μM, water | ryanodine receptors, L-type Ca<sup>2+</sup> channels | No effect |
| Ruthenium Red | Ammoniated ruthenium oxychloride | 11103-72-3 | 50 μM, water | Transient receptor potential channels, ryanodine receptors, voltage-sensitive Ca<sup>2+</sup> channels, Ca<sup>2+</sup> ATPases, calmodulin | No effect |
| U-73122 | 1-[6-[[[(17β)-3-Methoxyestra-1,3,5(10)-trien-17-yl]amino]hexyl]-1H-pyrrole-2,5-dione | 112648-68-7 | 25 μM, DMSO | Phospholipase C | No effect |
| H-8 | *N*-[2-(Methylamino)ethyl]isoquinoline-5-sulfonamide dihydrochloride | 113276-94-1 | 10 μM, DMSO | Protein Kinases A, C, and G, myosin-light chain kinase | No effect |
| Methylene Blue | 3,7-bis(Dimethylamino)-phenothiazin-5-ium chloride | 61-73-4 | 30 μM, water | Nitric oxide synthase, guanylate cyclase | No effect |
| NS-2028 | 8-Bromo-1H,4H-[1,2,4]oxadiazolo[3,4-c][1,4]benzoxazin-1-one | 204326-43-2 | 10 μM, DMSO | guanylate cyclase (NO-induced) | No effect |
| QDQ | 1H-[1,2,4]oxadiazolo[4,3,-a]quinoxalin-1-one | 41443-28-1 | 10 μM, DMSO | guanylate cyclase (NO-induced) | No effect |

**Figure S1.**
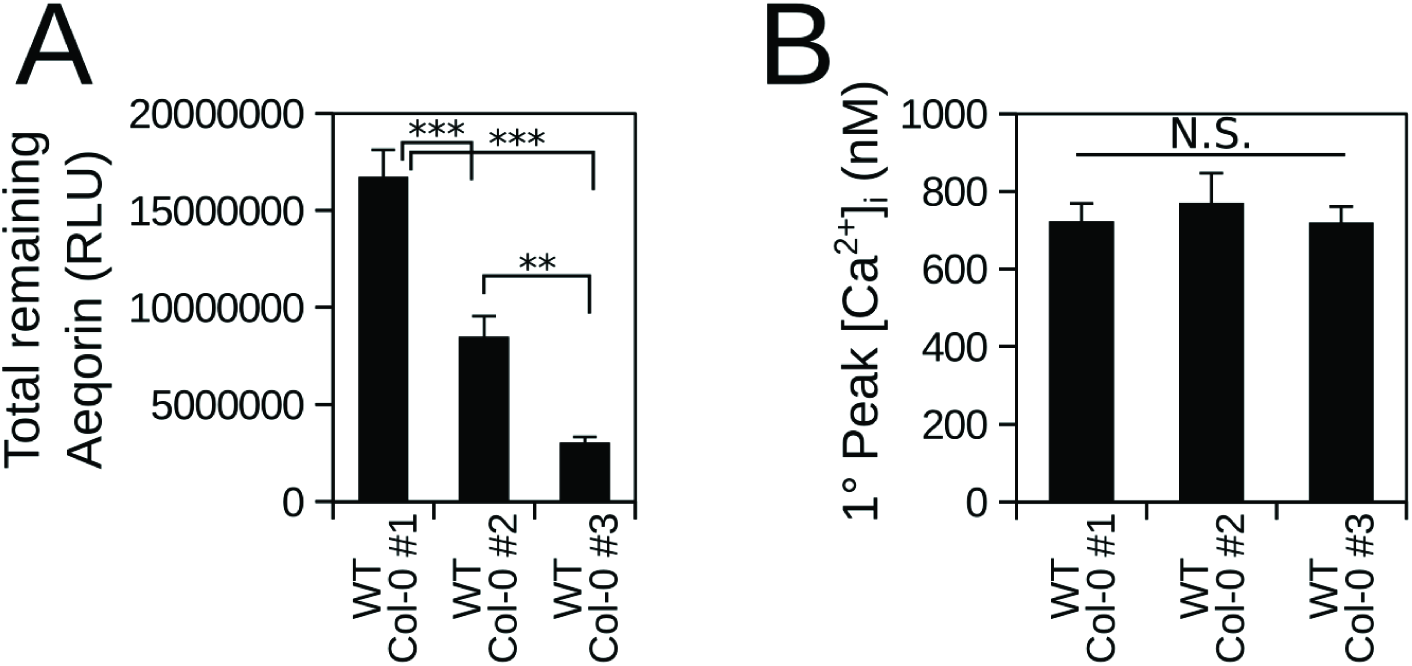
Calibrated Ca^2+^ measurements are independent of total aequorin expression levels. Three wild-type Col-0 lines independently-transformed with T-DNAs encoding the aequorin Ca^2+^ reporter under the control of the 35S promoter. (**A**) Average total counts of Aequorin remaining after hyperosmotic stimulation, determined by application of 2M CaCl_2_ + 20% ethanol. (**B**) Average calibrated 1° peak Ca^2+^ measurements ± SEM in response to 316 mOsmol/L stimulation. See Methods for computation of free Ca^2+^. Statistical comparisons between lines were made by one-way ANOVA with Tukey-HSD post-hoc test. *** p<0.001, ** p<0.01, N.S. p>0.05. *n* = 15-27 seedlings per line.

**Figure S2.**
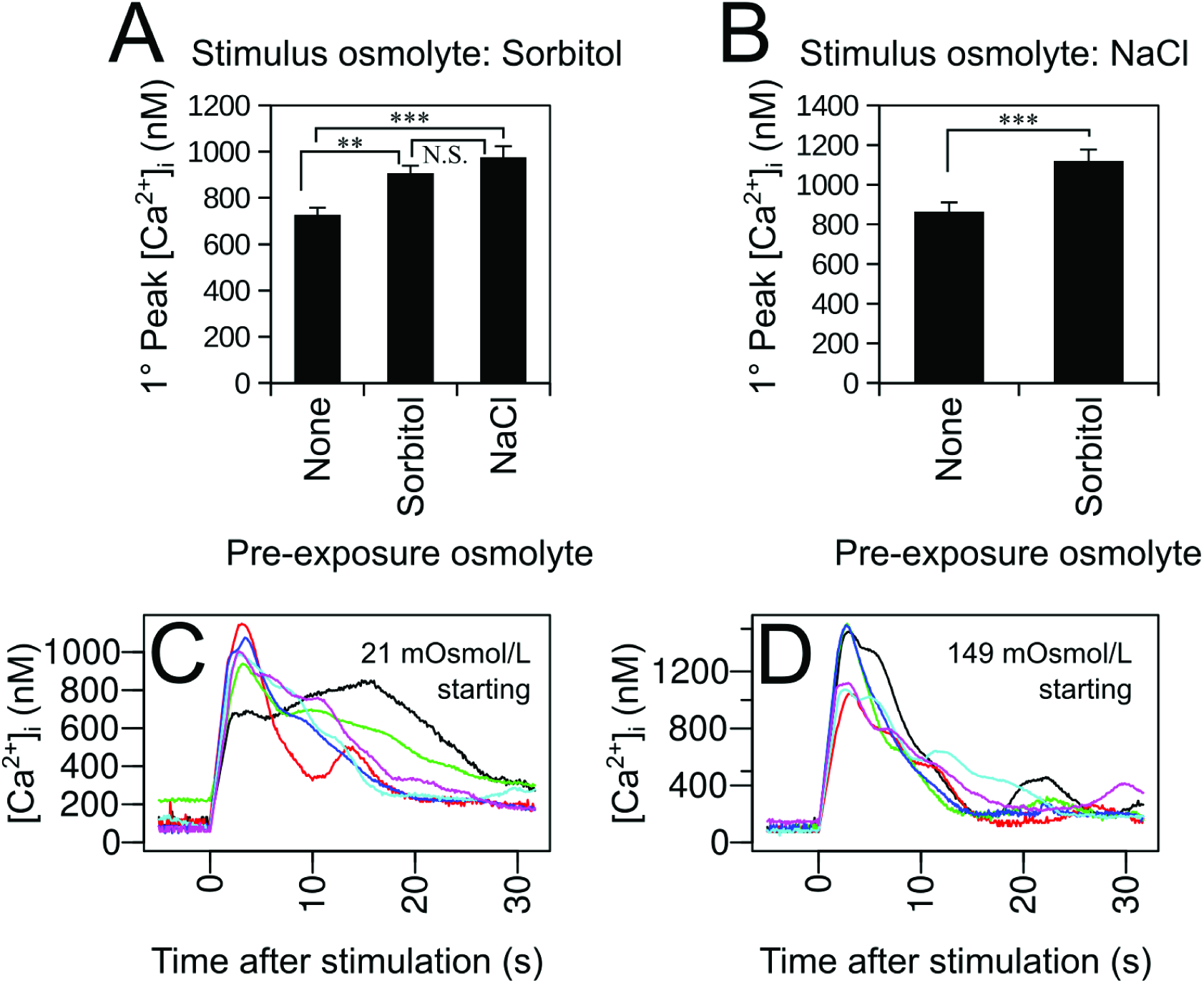
Cross-potentiation between sorbitol and NaCl stress and example individual responses. (**A**) Average 1° peak Ca^2+^ measurements ± SEM of seedlings stimulated with an injection of hyperosmotic sorbitol solution. The seedlings were pre-exposed to solutions of water (no additional osmolytes), or to moderate hyper-osmotic stress (149mOsmol/L) by application of sorbitol or NaCl solutions. *n* = 30-32 seedlings per treatment group. (**B**) Average 1° peak Ca^2+^ measurements ± SEM of seedlings stimulated with an injection of hyperosmotic NaCl solution. The seedlings were pre-exposed to solutions of water (no additional osmolytes), or to mild hyper-osmotic stress (149mOsmol/L) by application of sorbitol solution. Data in (**B**) are from a larger experiment that includes data from Figure 1A-G. Statistical comparisons between lines were made by one-way ANOVA with Tukey-HSD post-hoc test. *** p<0.001, ** p<0.01, N.S. p>0.05. *n* = 29 seedlings per pre-exposure group. (**C-D**) Six examples of individual seedling Ca^2+^ responses to 0.7 volumes of 700 mOsmol/L sorbitol starting from indicated solution osmolarity, resulting in final osmolarities of 316 (**C**) and 388 (**D**) mOsmol/L, respectively. Note the differing y-axis scales in C and D.

**Figure S3.**
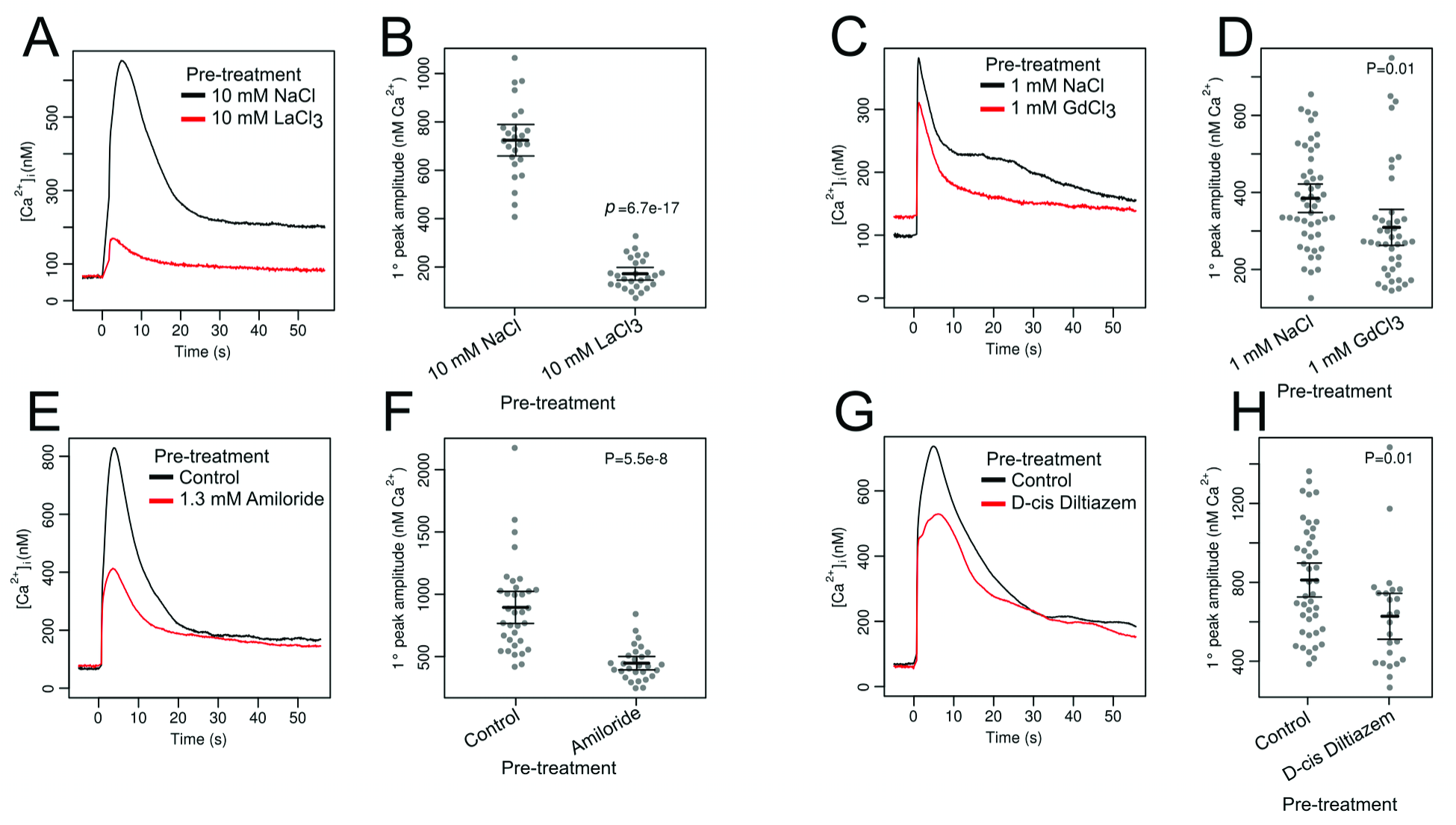
Pharmacological inhibition of rapid osmotic-induced Ca^2+^ responses. (**A-B**) Effects of 10 mM LaCh treatment. (**C-D**) Effects of 1 mM GdCl3 treatment. (**E-F**) Effects of 1.3 mM amiloride treatment.(**G-H**) Effects of 100 μM Diltiazem treatment. (A,C,E,G) Average Ca^2+^ response traces of 1-week-old whole seedlings stimulated with an injection of hyperosmotic NaCl solution at Time = 0. (B,D,F,H) Primary peak amplitudes. Each dot represents one seedling. Horizontal bars represent mean ± SEM. Student's T-test was used to calculate the *p*-values shown.

**Figure S4.**
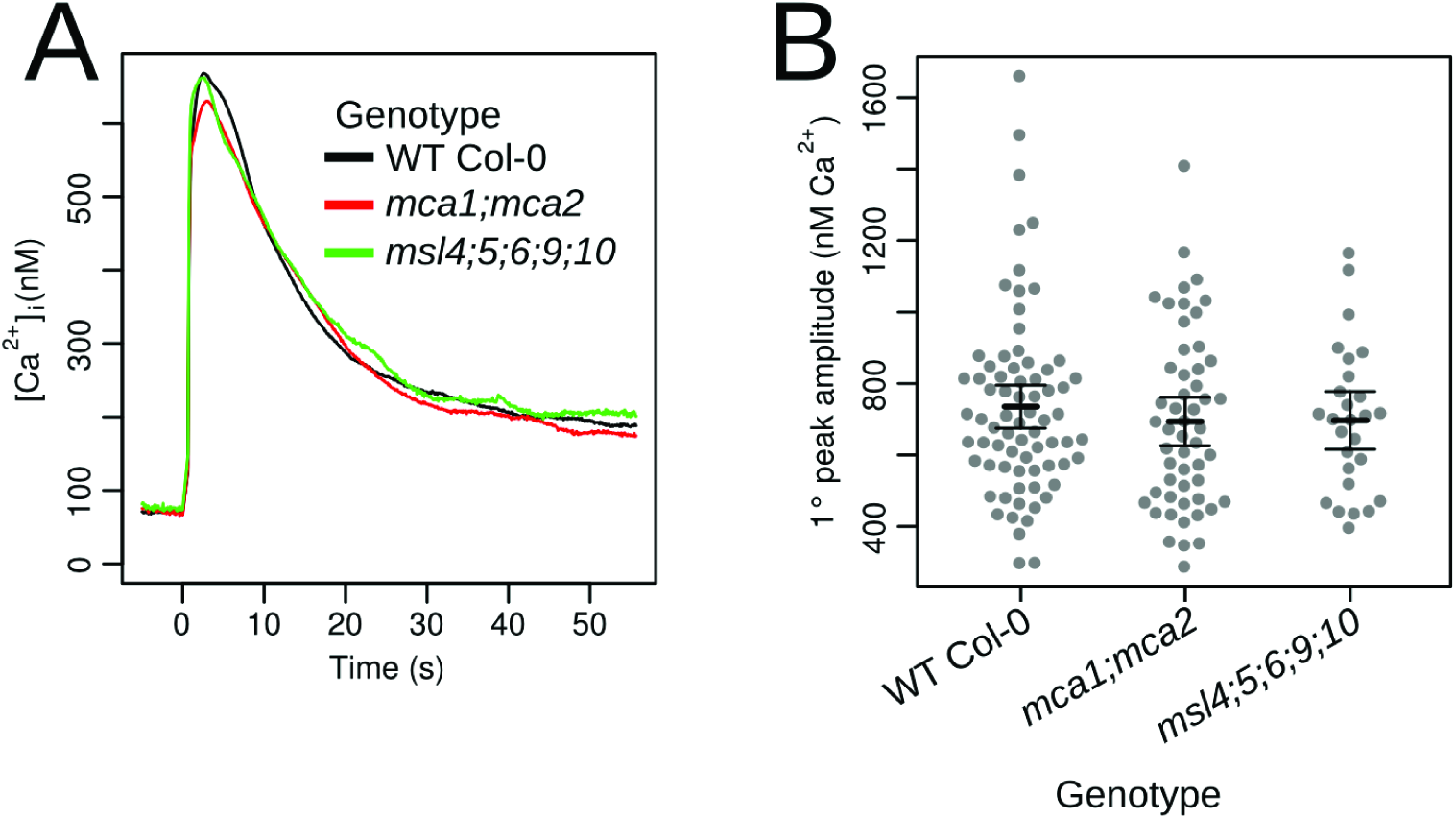
Analysis of osmotic-induced Ca^2^ responses in mechanosensitive channel mutants. (**A**) Average Ca^2+^ responses of 1-week-old whole seedlings stimulated with an injection of hyperosmotic NaCl solution at Time = 0. (**B**) Primary peak amplitudes. Error bars represent ± SEM. One-way ANOVA showed no significant differences between genotypes.

